# Easy and interactive taxonomic profiling with Metabuli App

**DOI:** 10.1101/2025.03.10.642298

**Authors:** SunJae Lee, Jaebeom Kim, Milot Mirdita, Cameron L.M. Gilchrist, Martin Steinegger

## Abstract

**Summary:** Accurate metagenomic taxonomic profiling is critical for understanding microbial communities. However, computational analysis often requires command-line proficiency and high-performance computing resources. To lower these barriers, we developed Metabuli App, an all-in-one desktop application that efficiently runs taxonomic profiling locally on a consumer-grade computer. It features user-friendly graphical interfaces for custom database curation, raw read quality control (QC), taxonomic profiling, and interactive result visualization.

**Availability and implementation:** GPLv3-licensed source code and prebuilt apps for Windows, macOS, and Linux are available at https://github.com/steineggerlab/Metabuli-App and are archived at https://doi.org/10.5281/zenodo.15876171. Analysis scripts are available at https://github.com/jaebeom-kim/metabuli-app-analysis. The Sankey-based taxonomy visualization component is available at https://github.com/steineggerlab/taxoview for easy integration into other web projects.

## Introduction

Metagenomic taxonomic classification allows researchers to understand the taxonomic structure of microbial communities by analyzing environmental DNA. However, it often requires command-line proficiency and high-performance computing resources.

Tools like MEGAN (1) and Pavian (2) provide graphical user interfaces (GUIs) for metagenomic analysis and visualization. However, MEGAN relies on alignment results, and Pavian requires taxonomic profiling results, both generated by separate tools that often demand server-level hardware and command-line skills.

Web applications like CZ ID (3), Galaxy (4), MG-RAST (5), and MGnify (6) lower technical barriers by offering GUIs and server-side computation. CZ ID provides pipelines for metagenomic analysis and pathogen detection. Galaxy offers diverse bioinformatics tools, including for metagenomics. MG-RAST and MGnify deliver comprehensive functional and taxonomic annotations.

However, they require registration, login, and data upload, which may be limited by data policies, storage quotas, or internet connectivity. For example, MGnify requires public data submission prior to analysis; the usegalaxy.org Galaxy server has a hard 250 GB total user quota, requiring users to use alternative Galaxy servers or a local installation for larger projects. Analyses may also be delayed by server queues, and none of them support custom reference databases.

Building on these prior efforts, we present Metabuli App—the first stand-alone desktop application supporting a full suite of functionalities: database download and curation, raw read QC, taxonomic profiling, and result visualization. Inspired by Pavian (2), the visualization features interactive and customizable Sankey (7) and Krona (8) plots. Except for database download, all processing is performed locally, eliminating the need for uploads, internet access, or remote servers.

The app includes a desktop-optimized version of Metabuli (9) as its classification engine. As benchmarked in Kim et al. (9), Metabuli achieves high classification performance by comparing query and reference sequences at both amino acid and DNA levels, integrating sensitive detection of novel species with precise differentiation of closely related taxa. By lowering computational barriers while supporting key capabilities, the app makes advanced metagenomic profiling more accessible to a broad range of users.

## Methods

### Metabuli App implementation

Metabuli App is built using Electron (electronjs.org) to support Windows, macOS, and Linux. The front-end utilizes Vue (vuejs.org) for an interactive GUI, with D3 (10) rendering Sankey plots and Krona rendering radial hierarchical plots. The app internally runs fastp v1.0.1, fastplong v0.3.0, and Metabuli v1.1.1 for short-read QC, long-read QC, and taxonomic classification, respectively, based on parameters set by the user through the GUI. As these tools were originally built for UNIX-like systems, we added Windows support via Cygwin, a POSIX layer enabling cross-platform compatibility. To facilitate this, we replaced the Makefile-based build systems of fastp and fastplong with CMake. Additionally, for fastplong, we replaced the default Cygwin memory allocator with nedmalloc from the Git for Windows project (11), resulting in over an eightfold speedup.

#### Sankey plot generation using Taxoview

Sankey plots intuitively display the distribution of taxa across taxonomic ranks. We developed a standalone Vue.js component for Sankey visualization, Taxoview (12), which we also recently integrated into the Foldseek search (13) and cluster (14; 15) webservers. It takes a taxonomy report in Kraken format (16), hierarchically extracts the rank, proportion, and clade read count for each taxon and internally converts the results into a JSON structure, then renders an interactive Sankey plot. The plot is validated by reconstructing the original taxonomy report from the JSON.

##### Filtering and customization

Users can set filtering thresholds for minimum proportion, clade count, and most abundant taxa per rank. Visualizations update dynamically as thresholds are adjusted.

##### Subtree Sankey plot

The user-specified node becomes the root, and all descendant nodes and links are recursively gathered, applying the same filtering criteria used in the main Sankey plot. The filtered set of nodes are then processed again by the Taxoview component to update the Sankey plot.

### New Metabuli commands

The `updateDB` command computes a sorted `k`-mer list from new sequences and merges it with that of an existing database, and identical `k`-mers from the same species are dedupli cated. The merged `k`-mer list becomes the updated database.

The `extract` command retrieves reads classified under a specified taxon or its descendants. It takes a taxonomy ID, scans the classification results (which retains read order), collects the line numbers of matching classifications, and extracts the reads corresponding to these numbers into a separate file.

### Metabuli optimization

#### Windows support and optimized memory use

Cygwin, a POSIX layer for Windows, introduced overhead particularly in memory-intensive and parallel operations, so we optimized memory use by replacing associative containers with linear ones and minimizing allocations, especially within parallel blocks. Additionally, the efficient IPS^4^o sorting algorithm (17), which was already enabled in Linux and macOS, was also activated for Windows.

#### Faster multi-threaded query *k*-mer extraction

Previously, a number of reads matching the thread count were loaded and distributed one per thread for `k`-mer extraction. Loading and `k`-mer extraction occurred sequentially, creating synchronization bottlenecks after each step.

Now, read loading and `k`-mer extraction occur concurrently with a granularity of 1,000 reads. The main thread continuously identifies an idle thread and loads 1,000 reads into its pre-allocated memory. Once a thread has its sequences assigned, it immediately begins extraction without waiting for the others.

#### Faster *k*-mer match assembly

When evaluating a species’ `k`-mer match list sorted by read coordinates, the main bottleneck was identifying and assembling consecutive matches—shifted by one codon with identical overlap—to maximize the score of the covered read region.

The previous approach built a directed graph with matches as nodes and edges linking consecutive ones. A depth-first search from each source node identified the highest-scoring path to a terminal node, whose coordinates and score were used to compute the species score. This method scanned the match list twice—once for graph construction and once for traversal.

We replaced it with a non-graph-based, cache-efficient algorithm that scans the match list only once First, matches at *n* and *n* + 1 are identified. Each match at *n* + 1 is connected to the highest-scoring consecutive match found at *n*. When connecting two matches, the preceding match’s score is added to the next match’s additional codon score to form the next match’s score, with the start position of the match chain carried forward. As the process advances, match chains emerge, extend, or terminate. Match chains are sorted by score to select the highest-scoring paths, the same ones identified by the previous approach.

#### Speed Measurement

Three systems were used: a Windows desktop (Intel i9-9900, 32 GB RAM), a MacBook notebook (M2 Pro, 32 GB RAM), and a Linux server (64-core AMD EPYC 7742 CPU, 1 TB RAM). Metabuli v1.1.1 was used with all available threads and `--max-ram` set to 32, 32, and 128 GiB on the respective systems, unless stated otherwise. Average time over five runs is reported, except for classification with v1.0.6 on Windows, measured only once due to long runtime.

For database creation, we used 8,520 genomes (33 GB) from the Genome Taxonomy Database (GTDB) Release 220 meeting the following criteria: species representative, Complete Genome or Chromosome-level assembly, CheckM2 completeness *>* 90%, and contamination *<* 5% (18). Scaffold-level assemblies meeting the same criteria were used in Fig. 1g. To update the GTDB database with viral genomes, we used the Master Species List 39.4 of International Committee on Taxonomy of Viruses (ICTV) (19), which includes 14,690 species, 14,558 of which have GenBank accessions. We used 18,672 accessions representing 13,596 species, retrieved from GenBank virus genomes downloaded in January 2025.

**Fig. 1.**
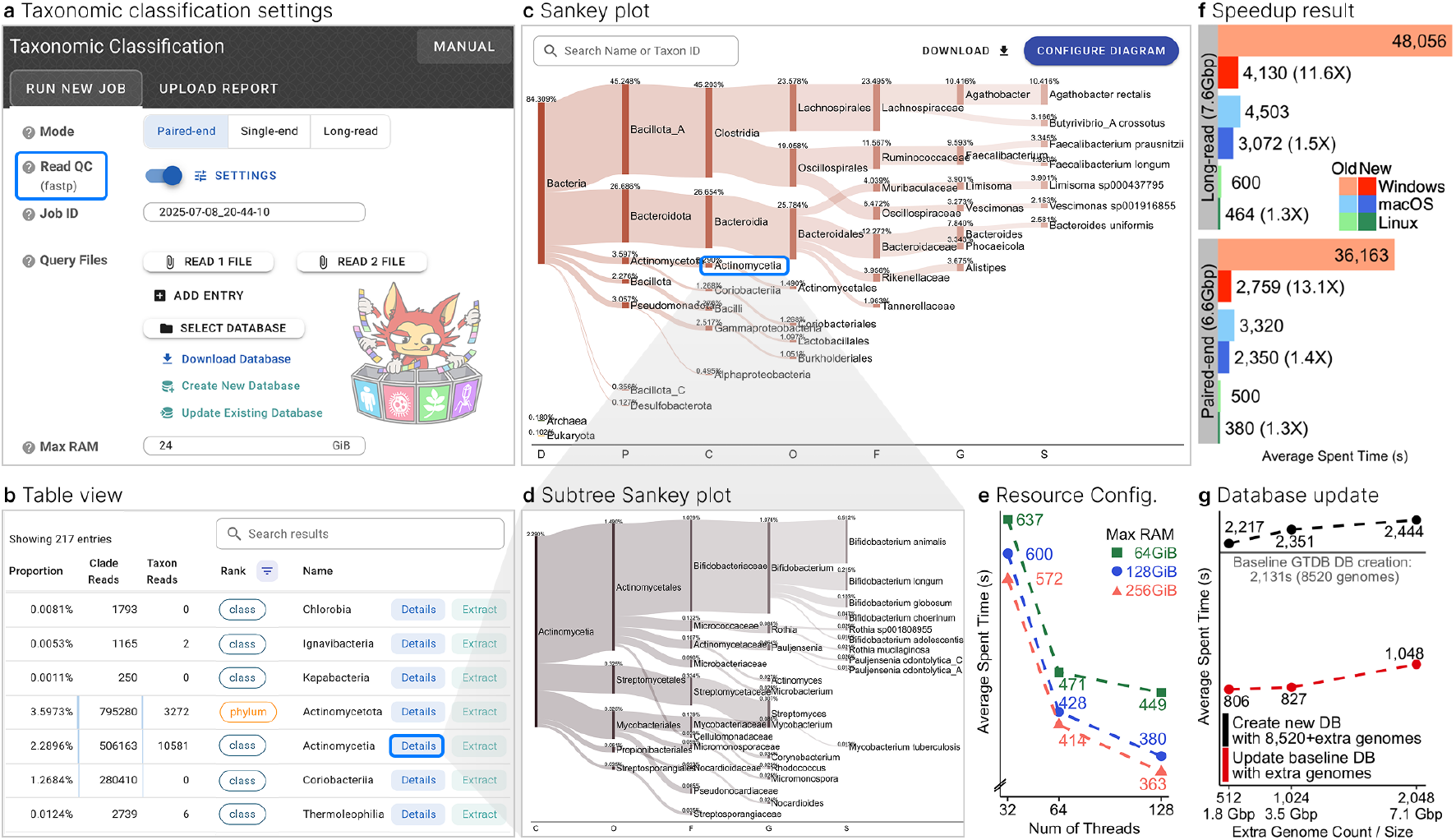
Metabuli App interface and Metabuli optimization results. **a**, Classification settings interface, with the raw-read QC highlighted. **b**, Classification summary table of detected taxa. Rank filtering displays only the phylum and class ranks. **c,d**, Sankey plots across the full taxonomic tree (**c**) and for a selected subtree (**d**). **e**, Classification time on a Linux server under varying resource settings. **f**, Classification speedup of Metabuli v1.1.1 (New) over v1.0.6 (Old) for paired-end (SRR24315757) and long-read (SRR15489014) human gut metagenomes. **g**, Time comparison between creating a new database and updating an existing one on the Linux server

## Features and Execution Times

### Database creation

In addition to downloading prebuilt databases, users can create custom databases against which reads are compared for taxonomic classification. This requires sequence files, taxonomy dump files, and an accession-to-taxonomy ID mapping. Metabuli supports Taxonkit(20)-generated dump files for GTDB (18) and ICTV (19), as well as NCBI Taxonomy (21). Creating a quality-filtered GTDB database took 36, 106, 143 minutes on a Linux server, MacBook, and Windows desktop, respectively.

### Database sequence addition

An existing database may lack certain clades of interest. In such cases, Metabuli now supports adding new genomes, which is faster than rebuilding the database from scratch (Fig. 1g) and does not require the original genome files. This allows users to easily expand prebuilt databases with additional clades. The taxonomy can also be extended using the `--new-taxa` option. Adding 13,596 viral species using ICTV taxonomy to the GTDB database created above took 13, 13, and 65 minutes on the same systems, respectively.

### Raw-read quality control

Raw-read QC, including quality filtering and adapter trimming, is performed by fastp for short reads and fastplong for long reads (22), both integrated into the app with adjustable parameters (Fig. 1a). Run times were measured using default parameters, except for using ten threads. A paired-end human gut metagenome (SRR24315757, 6.6 Gbp) took 39, 47, and 55 seconds on the Linux server, MacBook, and Windows desktop, respectively, while a long-read sample (SRR15489014, 7.6 Gbp) required about 36, 69, and 101 seconds.

### Taxonomic classification

After database preparation and raw-read QC, users can proceed to taxonomic classification. The app supports single-end, paired-end, and long-read FASTA/Q files (including gzip-compressed) and automatically validates their format (23; 24). Specifying a batch of read files, a database, and an output directory is sufficient to start the analysis. The output includes per-read classifications, a Kraken-format report (16), and a Krona HTML file (8). The app uses all available threads and memory by default, but this is user-configurable, affecting only runtime (Fig. 1e), not classification results. Users can also set a minimum score threshold to reduce false positives, at the cost of search sensitivity (9).

We measured classification speed using a database containing 36,203 genomes across 8,465 prokaryotic species (25; 26). Classifying the paired-end human gut metagenome took approximately 6, 39, and 46 minutes on the Linux server, MacBook, and Windows desktop, respectively, while the long-read human gut data required about 7, 51, and 69 minutes, respectively (Fig. 1f). This corresponds to approximately 1.3*×*, 1.4*×*, and 13*×* speedups for paired-end data, and 1.3*×*, 1.5*×*, and 12*×* speedups for long-read data over the previous version, while producing identical results.

In addition, classification scales linearly, enabling efficient analysis of larger datasets; the original, doubled, and quadrupled paired-end data took 39, 79, and 157 minutes on the MacBook. Moreover, Metabuli showed identical speed on the MacBook with both the database and paired-end reads stored on an external SSD (CORSAIR EX400U), demonstrating flexible storage options. For soil metagenomes, classification took 10, 67, and 87 minutes for paired-end data (SRR26074284, 6.6 Gbp), and 10, 66, and 89 minutes for long-read data (SRR13608742, 10.4 Gbp) on the Linux server, MacBook, and Windows desktop, respectively.

### Customizable and interactive result visualization

The classification results are visualized as a table, a Sankey plot, and a Krona chart. A previous report file can also be opened in the app to generate the table and Sankey plot.

#### Table view

It displays the read proportion and count for each detected taxon (Fig. 1b). The results can be filtered by taxonomic rank, sorted by column, and searched by taxon. The “Details” button opens the “Detailed View” of the taxon.

#### Sankey plot

It intuitively visualizes the distribution of classifications across the taxonomic hierarchy (Fig. 1c). Users can hover over nodes to highlight lineages, search for taxa, and access a taxon’s “Detailed View” by clicking its node. Customization options includes filtering low-abundance taxa, limiting taxa per rank, selecting rank to display, and adjusting visuals. The plot is exportable in PNG or SVG format.

#### Detailed view

It shows the full lineage and a customizable subtree Sankey plot of the selected taxon (Fig. 1d). When an NCBI taxonomy-based database is used, links to the corresponding NCBI taxonomy and genome web pages assist downstream analysis.

### Taxon-specific read extraction

The “Extract” button in the table and detailed view lets users collect reads assigned to a specific taxon into a separate file.

## Conclusion

Metabuli App addresses key barriers in taxonomic profiling: the need for server-level computing resources and command-line expertise, while also providing a full suite of functionalities, including database curation, raw-read QC, taxonomic classification, interactive visualization, and read extraction for taxa of interest. The Sankey plot component, Taxoview, offers flexible customization for publication-quality figures and can be easily integrated into other projects. Moreover, Metabuli App’s local processing ensures data privacy and enables use in low-connectivity settings such as remote clinics or field sites, and efficient performance on consumer-grade hardware reduces costs and broadens accessibility. Its intuitive interface helps users understand metagenomic workflows, making it a valuable educational tool, still supporting a full suite of advanced, customizable analysis functions. We anticipate that Metabuli App will enable a wide range of metagenomics researchers and practitioners, including those in microbiome research and clinical pathogen detection, to perform advanced taxonomic profiling without requiring command-line proficiency or specialized computing infrastructure.

## Funding

This work was supported by National Research Foundation of Korea [2020M3-A9G7-103933, 2021-R1C1-C102065, 2021-M3A9-I4021220, RS-2023-00250470, RS-2024-00396026]; Novo Nordisk Foundation [NNF24SA0092560]; Seoul National University [Creative-Pioneering Researchers Program, AI-Bio Research Grant]; and Samsung DS research fund. This work was supported by a grant from the National Institute of Biological Resources (NIBR), funded by the Ministry of Environment (MOE) of the Republic of Korea (NIBRE202505).

M.M. acknowledges support from the National Research Foundation of Korea (grant RS-2023-00250470). C.L.M.G. acknowledges support by the Ministry of Education of the Republic of Korea and the National Research Foundation of Korea (C07029) and the KBSI internal research programs (C539110).

## Author contributions

All authors contributed to the conceptualization of the application. S.L. developed the graphical user interface under the guidance of M.M., with contributions from C.L.M.G. to improve visualizations. J.K. optimized Metabuli’s performance. S.L. and J.K. conducted speedup measurements and, along with M.M. and M.S., co-authored the manuscript.

## Competing interests

M.S. acknowledges outside interest in Stylus Medicine. The remaining authors declare no competing interests.

## Notes

https://github.com/steineggerlab/Metabuli-App

https://github.com/steineggerlab/taxoview

### Summary of Updates

App updates (v1.1.1): Added multi-sample processing with fastp/fastplong QC, improved documentation, enhanced the Sankey visualization for publication-ready figures, and boosted Windows performance. Manuscript changes: Described these enhancements; expanded comparisons with MG-RAST, Galaxy, and MGnify; condensed Methods to emphasize Results; added new benchmarks of classification times across thread/RAM setups; and strengthened Conclusions with practical use cases.

